# Spontaneous network transitions predict somatosensory perception

**DOI:** 10.1101/2023.10.19.563130

**Authors:** Abhinav Sharma, Joachim Lange, Diego Vidaurre, Esther Florin

## Abstract

Sensory perception is essential for transforming incoming information in the brain into targeted behavior. Our brains are everlastingly active, and variations in perception are ubiquitously associated with human behavioral performance. Previous studies indicate that changes in spontaneous neural activity within local sensory areas correlate with the perception of ambiguous stimuli. However, the contribution of whole brain spontaneous networks to perception is not well understood. Using an ambiguous tactile temporal discrimination task, we demonstrate that the interaction between wholebrain networks in the seconds of the spontaneous pre-stimulus period also contributes to perception during the task. Transitions to a frontal and a multi-frequency network across the brain are essential for the correct percept. Conversely, incorrect percepts are mainly preceded by transitions to an alpha-parietal network. Brain transitions occur faster during the period before stimulus presentation for correct stimuli detection, suggesting the need for enhanced network flexibility during this phase.

**Significance statement:** Our brain is constantly engaged in processing sensory input and translating it into sensory perceptions. When confronted with ambiguous sensory information, individuals do not always have the same perceptual experience. We demonstrate that brain network transitions to frontal areas are essential for the correct percept. Conversely, incorrect percepts are mainly preceded by transitions to an alpha-parietal network. Correct stimuli detections are characterized by faster transitions, suggesting the need for enhanced network flexibility. These results extend our knowledge of perception by pointing to the relevance of whole-brain spontaneous networks and their dynamic properties.

## Introduction

Humans are not constantly engaged in tasks. But even at rest, our brains are buzzing with activity, not about any specific cognitive demand, called spontaneous activity. This spontaneous activity is not mere noise, as highlighted by the research on resting state networks.^1–4^ Rather, spontaneous activity might influence our perception and decisions. Compare your daily morning routine over two days. Even if everything remains the same, you will produce variability (good or bad) across tasks on different days. To understand such variability, it is important to understand the relevance of spontaneous neural fluctuations.

There is already some knowledge about fluctuations within local brain areas during the pre-stimulus period. Perception correlates with changes in neural activity within brain regions processing sensory information milliseconds before a task begins.^5–11^ Since local brain areas do not work by themselves, this knowledge is likely only the tip of the iceberg. Rather, whole brain networks are likely to be at play and influence the activity in local areas. Time-resolved techniques robustly identifying whole-brain networks from spontaneous data indicate that these brain networks at rest fluctuate and reconfigure at fast (subsecond) time scales.^12,13^ Therefore, differences in spontaneous network configurations and their temporal interplay during an extended period before a stimulus presentation might be responsible for perception differences.

We investigated the relevance of whole-brain networks during the extended pre-stimulus phase for tactile perception using an ambiguous tactile temporal discrimination task in the Magneto-encephalogram (MEG). Participants received two short consecutive electric tactile stimuli. They indicated with a button press the percept of either one or two pulses (Figure 1). Short inter-stimulus intervals are used to induce perceptual ambiguity because they cause the perception of only one stimulus despite two being applied. Using a staircase procedure, we determined the critical stimulus onset asynchrony (SOA) for each participant, i.e., the inter-stimulus interval inducing 50% correct detections of two stimuli. Applying a data-driven approach, the Hidden Markov Model ^12,14^, on the seconds of electrophysiological spontaneous brain activity before trials with ambiguous percept, we identified whole-brain network properties and interactions that predict each individual stimulus percept. We hypothesize that correct and incorrect detection involve different temporal configurations of the pre-trial whole-brain networks. Previous studies investigating tactile processing attribute an important role to the fronto-parietal cortices in complex tactile perceptual decision-making (for a review, see Romo and Rossi-Pool (2020)^15^). Hence, we focus on how fronto-parietal connections with brain regions primarily involved in tactile perception, i.e., sensorimotor cortices, and their temporal evolution cause trial-by-trial variability in tactile perception.

**Figure 1:**
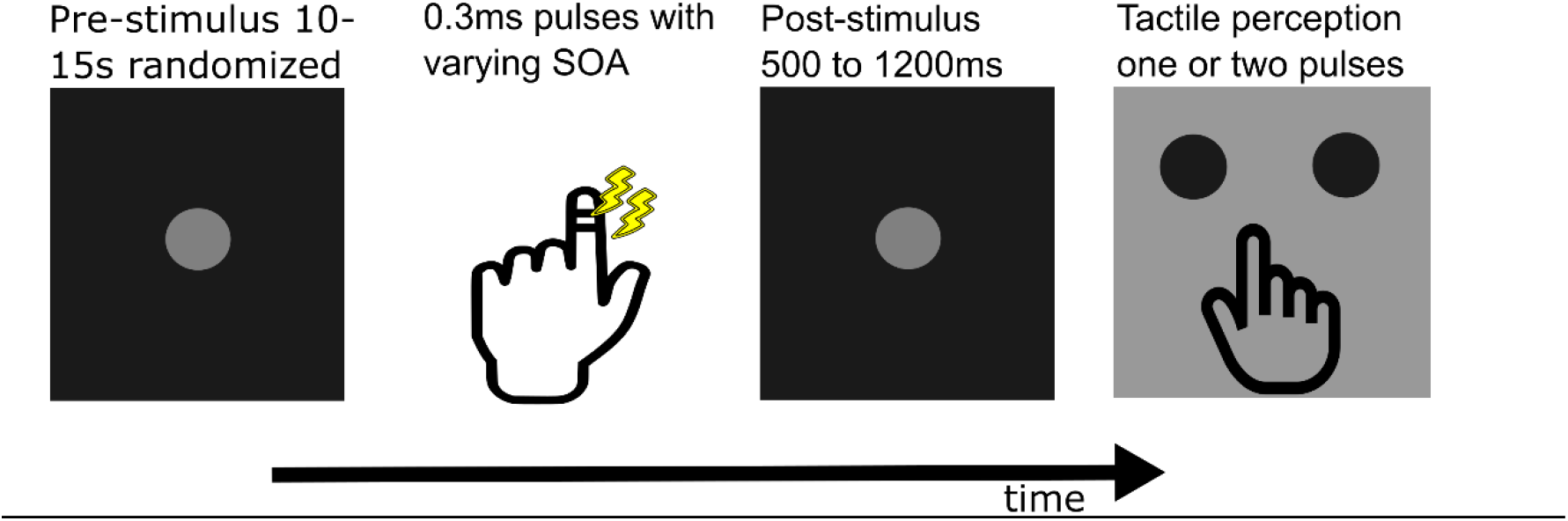
Schematic of the task paradigm (see Methods: Tactile discrimination paradigm), which was adapted based on previous studies (5,20). The presented stimulus onset asynchrony (SOA) varied between 0ms, the critical SOA, critical SOA ± 10 ms, and 3 times the critical SOA.

Our findings show two distinct whole-brain transition patterns related to correct and incorrect percepts, demonstrating the role of the interconnection and temporal interplay of spontaneous network configurations across the whole brain during the extended period before a task. This finding indicates that the literature on the neural mechanisms of human perception needs to include the role of dynamic whole-brain network patterns before a task.

## Results

Using a time-delay embedded (TDE)-Hidden Markov Model (HMM), we analyzed the spontaneous formation of whole-brain neural networks. Within the 8 seconds before the tactile stimulation, we identified 6 distinct networks. These networks were characterized by their spatial distribution and spectral fingerprint. Crucially, we found that the brain spontaneously transitions between these networks and that the network active at the moment of stimulation correlates with participants’ perception. In the first step, we will spatially and spectrally characterize the static network properties. These properties are, in turn, relevant for interpreting the dynamic properties of the networks associated with differing percepts.

### Spontaneous pre-stimulus activity organizes into distinct spectral networks

We identified six whole-brain networks with an HMM from the pre-stimulus time period of all critical SOA and critical SOA +-10ms trials regardless of the percept. We defined 42 cortical regions based on the Mindboggle atlas for further spectral and connectivity characterization. For the spectral characterization, we determined data-driven main frequency modes (see supplementary figure 1 and supplementary methods), which entail the alpha, beta, and high-beta/gamma bands. In contrast to canonical bands, these frequency modes allow for subject-specific weights of each frequency bin within a band.

First, we found one network that spans the whole brain, including frontal, temporal, and parietal areas (Fig. 2A, left part). This network was characterized by connections in multiple frequency bands, i.e., the alpha, beta, and high-beta/gamma bands. Interestingly, while alpha and beta-band connectivity is found across virtually all regions, the connectivity is sparser in the high-beta gamma band. Due to the contribution of all cortical areas, we termed this network *cross-brain* network. The second network was dominated by connections within and from the parietal cortex (Figure 2b). This connectivity was only present in the alpha band. Due to its spatial-spectral characteristics, we termed it the *alpha-parietal network*. The third network was dominated by spectral coherence in the high-beta/gamma and alpha-band in the frontal and medial-frontal regions and thus termed *frontal network* (Figure 2c). The fourth network was characterized by high-beta/gamma band coherence between and within frontal and parietal areas and termed *high-beta/gamma network* (Figure 2d). Importantly, the involvement of sensorimotor areas in each of these 4 networks suggests that they are indeed task-relevant. Finally, there were two networks with very sparse connectivity, thus putatively irrelevant for tactile perception (see supplementary figure 3). In the following, we focus on the four main networks.

**Figure 2:**
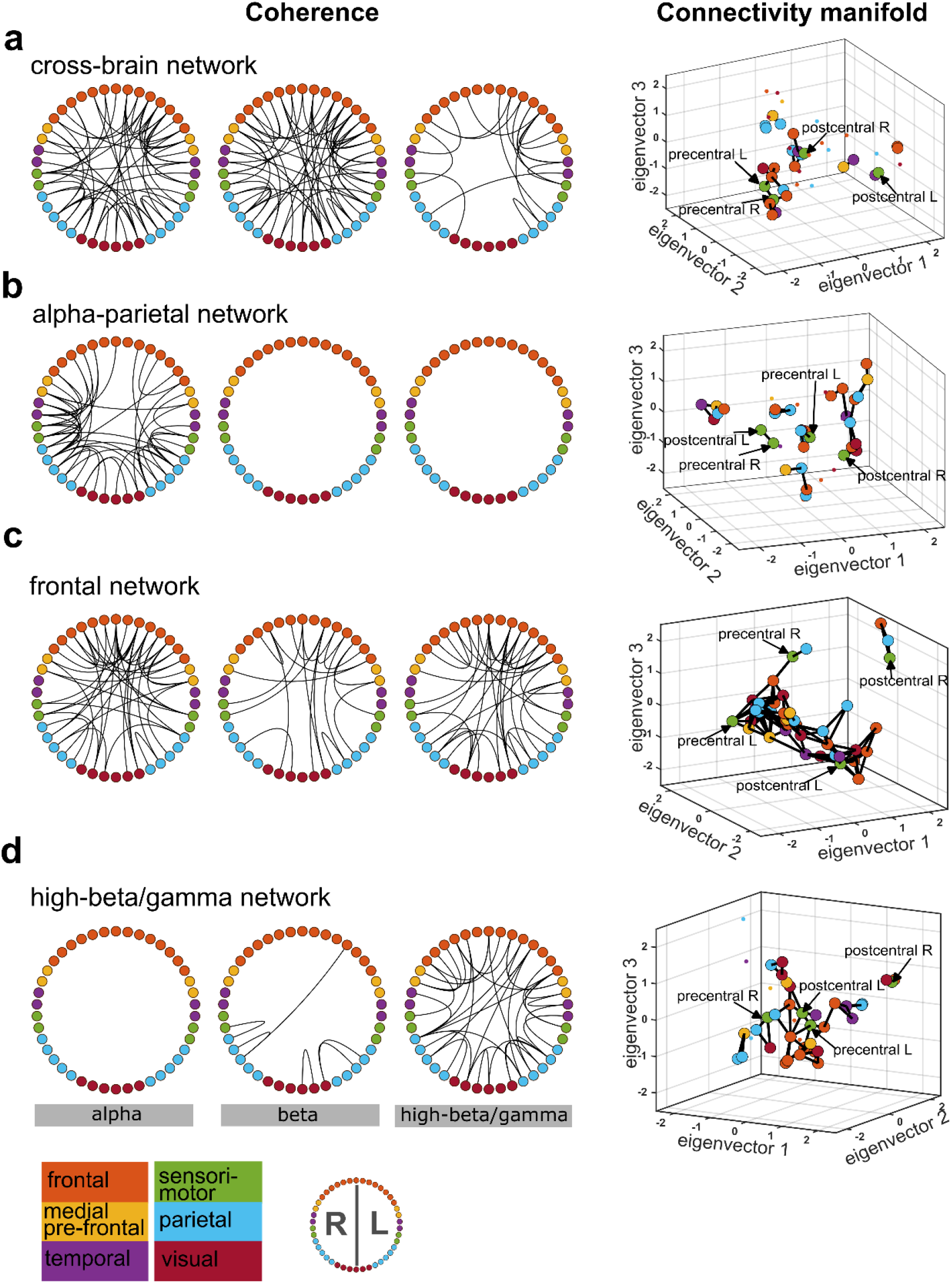
Coherence and connectivity of the four main networks. Each dot in the coherence ring and the connectivity manifold represents one of 42 cortical areas and is color-coded for the main brain region. The exact cortical areas are depicted in supplementary figure 2. **a)** *Cross-brain network*. The connectivity manifold for the cross-brain network indicated connections of the left and right precentral (motor cortices) with the frontal and parietal brain regions. **b)** *Alpha-parietal network*. This network was characterized by posterior alpha band coherence, but parietal cortex node connectivity was fragmented into different clusters. **c)** *Frontal network*. Frontal regions were coherent both within frontal cortex regions and to different brain regions across all three frequency bands. This network is densely connected within one main cluster. **d)** *High-beta/gamma network*. Significant connections were only present in the high-beta/gamma band both within the frontal and parietal regions as well as between the regions. The frontal and parietal regions form a cluster with left post-(somatosensory) and pre-(motor) central gyri.

### Connectivity clusters in the HMM networks

A brain region might operate simultaneously in different frequencies to perform multiple functions, enabling flexibility in the region’s inputs and outputs.^16,17^ Our previous section’s result indicated that numerous pairs of brain regions are coherent across several frequency modes within the same network. However, certain cortical regions were coherent in identical spectral bands across different networks. We calculated diffusion maps of each network to untangle whether this is because coherence cannot distinguish if a connection between two brain areas is direct or indirect, i.e., mediated via multiple brain areas.^18,19^ Diffusion maps extract a low-dimensional representation of data while preserving similarity calculated in higher dimensions. Figure 2 visualizes the maps on a three-dimensional connectivity manifold, which accounts for direct and indirect connections and places regions with greater connectivity closer to each other.

The right column of Figure 2 shows the connectivity manifold of each network. First, for the cross-brain network, only a few brain regions were statistically significantly connected with their neighbors on the cross-brain network’s connectivity manifold, indicating an integrative role of this network. This contrasts with the network’s multi-frequency and spatially non-specific coherence characteristics. Furthermore, the manifold shows that the parietal and frontal cortical areas, left motor cortex, and medial orbitofrontal regions were sparsely connected to each other (Figure 2a).

Second, the alpha-parietal network’s connectivity manifold displayed several disconnected clusters involving at least one parietal region. The parietal regions were, however, not connected to the left post-central and right pre-central gyrus (Figure 2b).

Third, all brain regions within the frontal network were connected in a single cluster, indicating a distributive role of the frontal network. This stands in sharp contrast to the three other networks. The only exception is the right post-central gyrus, forming a separate cluster with the superior-frontal and inferior-parietal right gyri (IP-r) (Figure 2c). Of note, the left and right pre- and post-central gyri were not connected in the frontal network.

Finally, the high-beta/gamma network showed a significant connection between the left pre-central and post-central gyri (Figure 2d and supplementary figure 6b for a 2D view of those connections). Like the cross-brain network, the IP-r was connected to the left pre- and post-central gyri via two frontal areas. The left pre- and post-central gyri were further embedded within a cluster of significantly connected frontal regions.

To summarize, the four HMM brain networks display distinct connectivity profiles between the sensorimotor cortices and frontal and parietal areas.

### Correct and incorrect detection correlates with different network transition probabilities

We need to rely on dynamic network analysis to test our hypothesis that correct and incorrect stimuli detections involve different temporal configurations of the whole-brain networks during the pre-stimulus period. The HMM analysis is ideally suited for this purpose as it naturally provides dynamic network features like transition probabilities and the duration spent in each network. To investigate our hypothesis, we examined on a single trial level whether transition probabilities to and from a given brain network significantly differed between correct and incorrect percept trials during the pre-stimulus period. We test for significantly different transition probabilities by testing which transition probabilities are significantly increased in each percept relative to the other (implying they are decreased in the other condition).

For trials in which participants incorrectly perceived one stimulus, the transition probability during the 8-second pre-stimulus period was higher from the frontal network to the alpha-parietal and the high-beta/gamma network (Fig. 3a, see supplementary figure 5 for the transitions to and from networks 5 and 6). In contrast, the likelihood of correct stimuli detection increased when the transition probability was higher from the alpha-parietal network to the cross-brain, the frontal, and the high-beta gamma network, as well as from the high-beta/gamma network to the cross-brain, the frontal, and the alpha-parietal network (Fig. 3b). The high-beta/gamma and alpha-parietal networks for correctly detected stimuli were the only pair with an increased transition probability in both directions.

**Figure 3:**
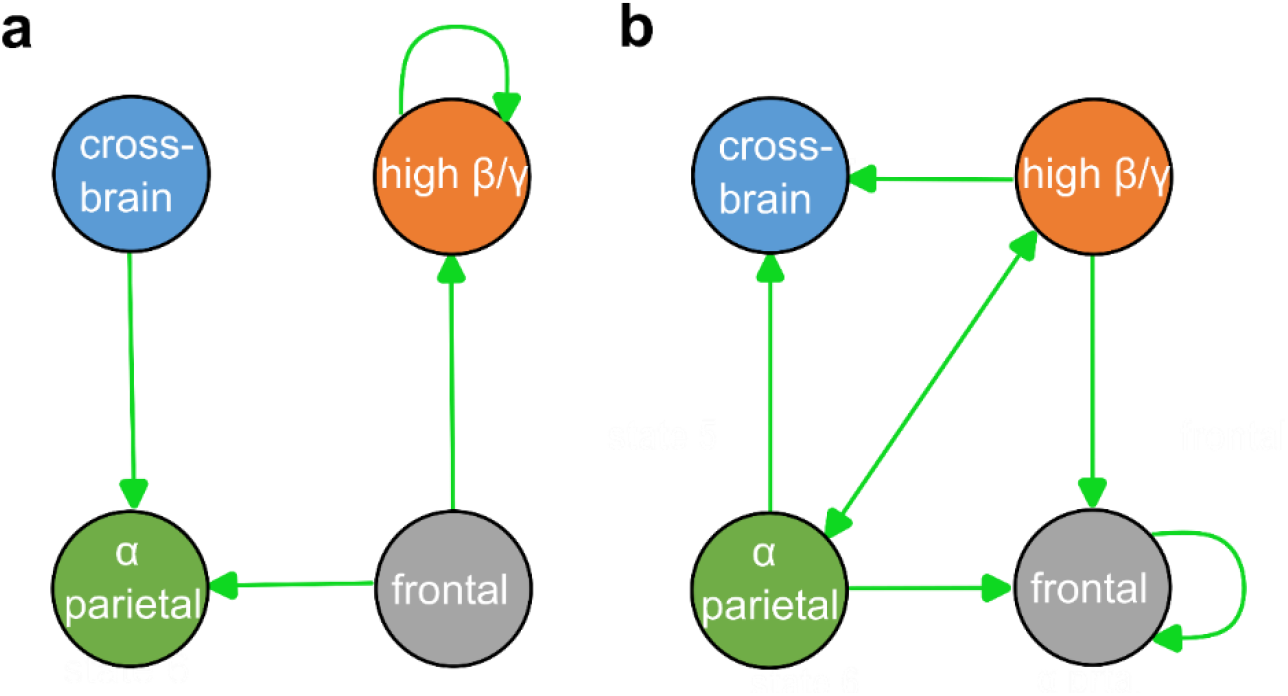
Pre-stimuli network transition probabilities differing across percepts. The arrows indicate significantly increased network transition probabilities between networks at p<0.05. Not depicted transition probabilities were statistically indistinguishable across percepts. **a)** Significantly increased network transition probabilities when participants incorrectly detected only one stimulus. **b)** Significantly increased network transition probabilities when participants correctly detected two stimuli.

### Faster network transitions improve tactile perception

Whether one or two stimuli were perceived crucially depended on the network transitions preceding them. We now focus on three dynamic characteristics of network transitions and investigate their role in accounting for differences in percepts across trials. First, we assess the temporal presence of networks, both in terms of the 1) duration of each visit (lifetime of a network) as well as 2) the overall share of time spent in a particular network (fractional occupancy (FO)). Second, we consider the rate of inter-network transitioning, measured by the number of network visits per second during the pre-stimulus period. Results are shown in Figure 4.

**Figure 4:**
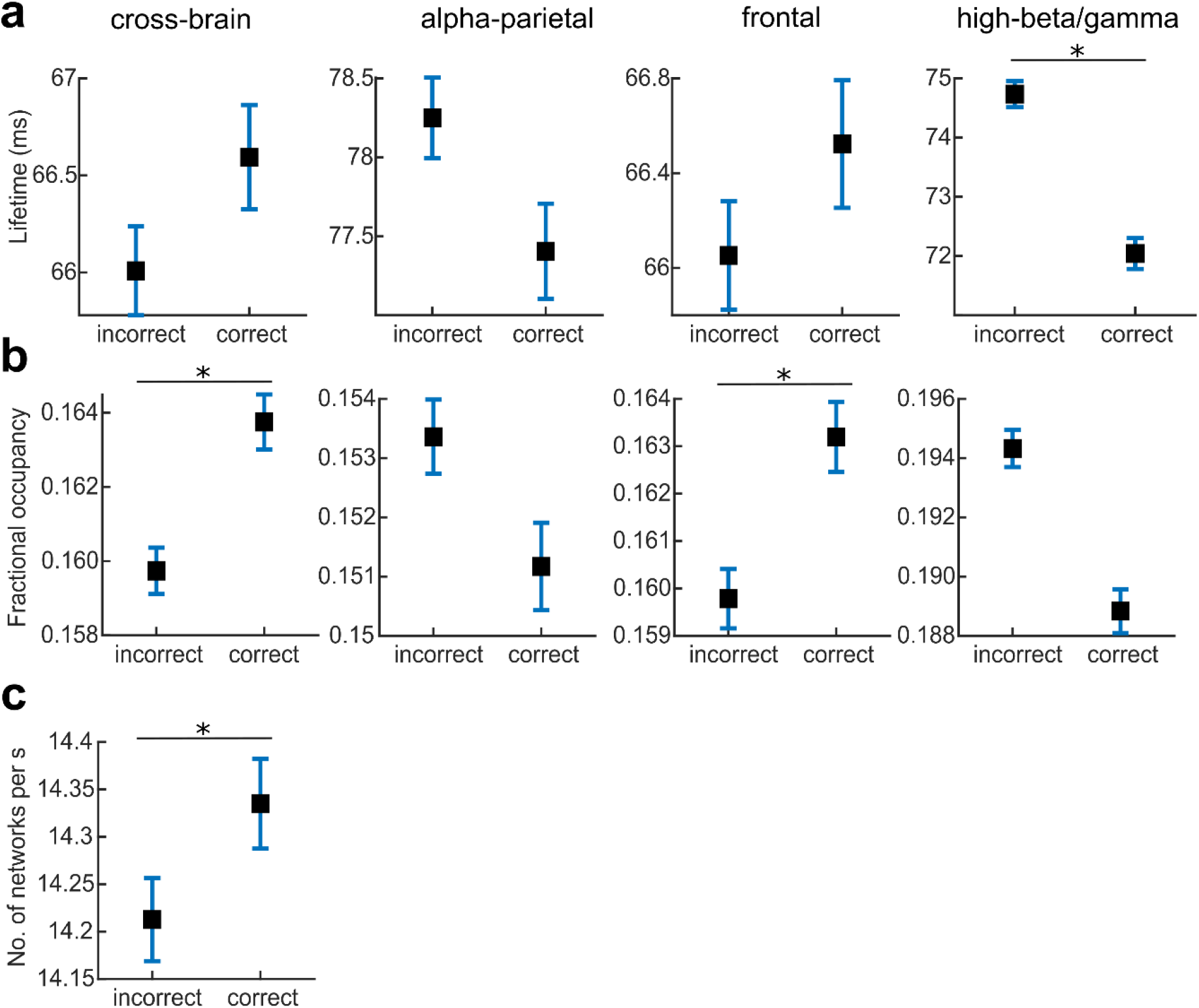
Temporal network properties. *Incorrect* denotes incorrect stimuli detection trials and *Correct* the correct stimuli detection ones. Box-whiskers represent the mean and standard error across all trials and participants. The stars denote statistically significant results with p<0.01. **a)** Lifetime for the corresponding HMM network **b)** Fractional occupancy (FO) for the corresponding HMM network **c)** Number of network visits per second for all six networks The lifetime and FO of the 2 further networks are presented in supplementary figure 6.

The lifetime of the alpha-parietal network was the highest (alpha-parietal > all other networks; p<0.001), and its lifetime did not significantly differ between correct and incorrect detections (p=0.59; Fig. 4a). For the high-beta/gamma network, the lifetime and FO were significantly lower for correct than incorrect detections (p<0.001). The cross-brain and the frontal network had a higher FO for correct than incorrect stimulus detections (p<0.001; Fig. 4b). In addition, correct stimulus detections were accompanied by faster overall inter-network transitions as the number of network visits per second was significantly increased.

## Discussion

Correctly detecting incoming stimuli in an uncertain, non-cued environment requires a particular dynamic pattern of pre-stimulus spontaneous brain activity that allows the brain to integrate the incoming stimuli optimally. In our work, we first identified four whole-brain networks relevant to the percept during the pre-stimulus resting period. Not all of those networks were simultaneously active, but during the pre-stimulus period, these networks fluctuated and reconfigured in a percept-dependent manner.

Correct percepts were characterized by higher transition probabilities from the alpha-parietal and high-beta/gamma networks to the frontal network and lower probabilities to the cross-brain network. In addition, the temporal characteristics of the transitions were relevant for the percept: Faster inter-network transitions preceded correct stimulus detection, suggesting a need for higher flexibility in the brain.

### Multi-frequency networks for correct percepts

The two detected multi-frequency networks, i.e., the frontal and cross-brain network, either distribute information to spectrally-specific networks in the case of the frontal network or assimilate information from such networks in the case of the cross-brain network. To process these in- and outputs, the cross-brain and frontal network require multi-frequency coherence to synchronize information across multiple frequencies. The cross-brain network is integrative because when higher-order brain interactions are considered, the connectivity is restricted to the left and right pre-central gyri, frontal, and parietal regions. In contrast, the frontal network is distributive because multi-frequency coherence is accompanied by a densely connected manifold. This fits well with the notation that frontal cortical regions perform heterogeneous and complex functions via multiplexed oscillatory coding.^17,20–24^ Hence, the frontal network is plausibly responsible for multiplexed oscillatory coding: the frontal cortex modulates coupling with other brain areas, allowing information to flow through the whole brain. This is achieved through dense connections to other brain regions.

### Spectrally-specific networks in relation to known power changes

In the case of the high-beta/gamma network – a spectrally-specific network – the left post- and precentral regions were embedded within prefrontal areas. Of note is the right IP-r’s presence in the prefrontal and left sensory-motor regions’ cluster. The static spectral properties and the dynamic connectivity properties for the high-beta gamma network align with previously reported findings in literature: In the sensorimotor system, beta activity is considered inhibitory^25,26^, while in the prefrontal regions, beta activity tends to indicate reactivation of neuronal ensembles for further processing.^27–29^ Moreover, it is known that IP-r in the human brain detects salient stimuli within sequences and controls attention over time.^30,31^ Hence, the high-beta/gamma network includes an ensemble of fronto-parietal cortical regions that activate perceptually relevant sensory-motor regions.

The spectral characteristics and the connectivity of the alpha-parietal network are consistent with the modulating role of parietal alpha activity for attention.^32–34^ Dense pair-wise alpha-band coherence between brain regions in the alpha-parietal network exhibited sparse connectivity dominated by parietal areas. The sparse parietal connectivity fits the described role of posterior alpha activity in gating cortical processing.^35–37^ Furthermore, the lifetime of the alpha-parietal network was the longest among all networks and did not significantly differ between correct and incorrect stimuli detections. Thus, the alpha-parietal network was not suppressed for correct stimuli detections, in contrast to the well-described task-related alpha power suppression in posterior areas.^32,35,36,38^ Instead, for correct stimuli detection, transitions occur back and forth between the alpha-parietal network and the high-beta/gamma network, and the alpha-parietal network has a higher probability of transitioning to multi-frequency networks.

### Connectivity manifolds for network descriptions

HMM is an unsupervised method that is blind to the underlying biology of the time series on which it is fit. Therefore, the extracted patterns are not biased by preconceptions or existing information about the underlying biology. Hence, the connection between biology and the HMM networks is necessarily post-hoc. At the same time, the HMM framework provides an estimate of the network dynamics through the lens of transition probabilities. This provides relevant information on network dynamics. However, these probabilities do not impose that a brain network is completely inactive. Rather it is an indication of dynamic network fluctuations and reconfigurations.

In our case, the cross-brain and the frontal network exhibited multi-frequency and spatially distributed spectral coherence. This overlap in coherence between the identified networks further complicates the interpretation of each network’s physiological relevance. Hence, we used diffusion maps on network-specific covariance vectors of each brain area to capture higher-order connectivity between different brain regions within each HMM network. The low-dimensional data representation of the diffusion maps allowed identifying a brain network’s most critical connections. Therefore, this approach enabled disentangling the integrative role of the cross-brain network and the distributive role of the frontal network by identifying a sparser connectivity profile. Altogether, diffusion maps offer a complementary framework to coherence-based whole-brain networks.

## Conclusion

Over the years, specific brain regions and their spectral properties have been linked to perception. However, brain areas do not operate in isolation but are intensely connected with each other. We highlighted that the likelihood of correctly detecting the tactile stimuli increased when transitions from spectrally-specific to multi-frequency networks were more likely during the pre-stimulus period. These changes in temporal network characteristics indicate that correct stimulus detection requires whole-brain networks to transition more flexibly and reconfigure network connectivity between sensorimotor, frontal, and parietal regions. Our findings collectively underscore the relevance of spontaneously forming whole-brain networks before the task for understanding mechanisms of perception. This extends previous research on near-threshold stimulus detection beyond the relevance of the short preparatory baseline period^8,9,39–41^ and thus points to the functional relevance of spontaneously forming whole-brain networks.

## Materials and methods

### Tactile discrimination paradigm

Thirty healthy participants (15 females and 15 males, right-handed, 26.4 (mean), 25 (median), 13 (range) years in age) volunteered to perform the task. The experimental procedure was explained to all participants. Everyone provided written consent, and the study was approved by the local ethics committee at the Medical Faculty of Heinrich Heine University, Düsseldorf (study number 2019-477) and conducted in accordance with the Declaration of Helsinki.

We adopted the tactile discrimination task described in detail in Baumgarten et al. (2016) to accommodate our network analysis. Each trial started with the presentation of a start cue (500 ms). Next, the cue decreased in brightness, indicating the pre-stimulus period (randomized to last between 10-15 seconds), after which the subjects received either one or two short (0.3 ms) electrical pulses applied by two electrodes placed between the two distal joints of the left index finger. The time between the two pulses (Stimulus Onset Asynchrony (SOA)) was determined with a staircase procedure for each participant prior to the main experiment so that they had a hit rate of 50%. This SOA is termed the *critical SOA* (crit SOA). Participants had to respond to whether they perceived one or two stimuli with their right hand by pressing a button with either their right index or middle finger. The index and middle finger options were randomly swapped at each trial. SOAs varied in a pseudorandom manner between 0ms, crit SOA, crit SOA + 10ms, crit SOA – 10ms, and 3 times crit SOA. The 0 and 3 times crit SOA trials were used to prevent biased responses and to confirm that the participants were properly performing the task. Participants were recorded on two days: Day one, for each participant, started with a pre-task, eyes-open, resting state MEG recording, which lasted for 10 minutes. It continued with the stair-case procedure and the MEG data recordings during the task. The task was divided into four blocks per day of about 45-50 trials. Each block lasted about 10-15 minutes. Participants were allowed to take breaks between blocks. Finally, a 10-minute post-task eyes-open resting state recording was performed.

On day two, the staircase procedure was repeated. This was followed by four measurement blocks of tasks during which MEG data were recorded. In total, across the two days, 200 trials with crit SOA, 50 of the other four conditions were acquired per participant. On the first day, crit SOA across subjects was 71ms (mean) +-34.3ms (standard deviation); on the second day, crit SOA across subjects was 79.8ms (mean) +-34.0 ms (standard deviation). The overall average crit SOA was 75ms (mean) +-34.0 ms (standard deviation). The accuracy across different SOAs is shown in supplementary figure 7.

### Electrophysiological recording

MEG data were recorded on both days of task performance. We used a whole-head MEG system with 306 channels (Elekta Vectorview, Elekta Neuromag, Finland) housed within a magnetically shielded chamber. Electrooculography and electrocardiography were recorded simultaneously with MEG data to later remove eye blink and cardiac artifacts. Data were acquired at a sampling rate of 2500Hz with an 833Hz low pass filter. The details of MEG-data pre-processing and source reconstruction can be found in the supplementary methods.

### TDE-HMM pipeline

The HMM is a data-driven probabilistic algorithm that finds recurrent network patterns in a multivariate time series^14,42^. Each network pattern is referred to as a ‘state’ in the HMM framework, such that these networks can activate or deactivate at various points in time. We use network throughout the text when referring to an HMM state. We used a specific variety of the HMM, the time delay embedded (TDE)-HMM, where whole-brain networks are defined in terms of spectral power and phase coupling^12^. For every point in time, the HMM algorithm provides the probability that a network is active. In our approach, we also performed spectral analyses of these networks, leading to a complete spatio-spectral connectivity profile across the cortex. A summary of the MEG data processing is provided in supplementary figure 8. Details on the HMM algorithm, the extraction of the data-driven frequency bands, coherence, and diffusion maps, can be found in the supplementary material.

### Group-level coherence ring figures

To separate background noise from the most robust coherent connections, a Gaussian mixture model (GMM) was used.^12^ For the group-level coherence projections, we normalized the activity in each network for each spectral band by subtracting the mean coherence within each frequency mode across all networks. As a prior for the mixture model, we used two single-dimensional Gaussian distributions with unit variance: one mixture component to capture noise and the other to capture the significant connections. This GMM with two mixtures was applied to the coherence values of each network. Connections were considered significant if their p-value after correction for multiple comparisons (Bonferroni correction) was smaller than 0.001.

### Statistical testing on the connectivity manifold

The connectivity manifold indicates the distance between different brain regions. In short, the smaller the distance between two regions on the manifold, the stronger the connectivity between the two regions (direct and indirect combined). For details about the relationship between distance and connectivity, please refer to Methods: Diffusion map analysis. To test if certain brain regions showed significant connectivity on the manifold, we applied the same test described in Methods: Group level coherence ring figures.

### Transition matrix test

We estimated transition matrices for each trial. For every trial, a transition matrix contains probabilities for different HMM networks to transition to and from one another. Non-parametric permutation testing (5000 permutations) was performed to compare network transitions between correct and incorrect trials. Results were considered significant if p<0.05 after correction for multiple comparisons using the Benjamini Hochberg FDR correction.^43^

### Temporal properties

To test for changes in the temporal properties for percept one versus percept two stimuli, we compared the lifetimes and FO for each network within and across the response types using two-way repeated measures ANOVA followed by post-hoc tests. A network’s lifetime for a given visit refers to the time spent in that network. Of note, the lifetime of a network can be shorter than one oscillation cycle because the frequency is the rate of change of a signal, and this rate can be estimated by examining the slope of the signal at any instant in time.^12^ FO is defined as the average share of time spent in a particular network. Finally, we assessed the overall network changes per second.

## Acknowledgments

This work was supported by the DFG grant FL 760/4-1 awarded to Esther Florin. We want to thank Lena Spitz for her assistance in recruiting participants for the study. We would also like to thank Johannes Pfeifer for his feedback on the manuscript.

## Author Contributions

AS: Conceptualization, Data acquisition and quality control, Software, Formal analysis, Investigation, Visualization, Methodology, Writing - original draft; JL: Behavioral paradigm, Methodology, Writing - review and editing; DV: Software, Writing - review and editing; EF: Conceptualization, Resources, Supervision, Funding acquisition, Validation, Investigation, Methodology, Visualisation, Project administration, Writing - review and editing

## Data and software availability

The raw data used in the study will be uploaded to a suitable data sharing repository post publication. The software developed is available on https://github.com/saltwater-tensor/tactstim_rest_publication_code.git.

