## Supplementary information for "Spontaneous network transitions predict somatosensory perception"

### Methods

#### MEG pre-processing

All data processing and analyses were performed using Matlab (version R 2021b; MathWorks, Natick, MA) and the Brainstorm toolbox[1]. The Neuromag system provides signal-space projection (SSP) vectors for cleaning environment noise from the MEG channels, which were applied. The line noise was removed from all channels with a notch filter at 50, 100, 150, ..., 550, and 600 Hz with a 3 dB bandwidth of 1 Hz. To ensure artifact-free data, we visually inspected the data. Very noisy and flat MEG channels were excluded from further analysis. Time segments containing artifacts were marked in the time series. However, if artifacts regularly occurred only in one channel, this whole channel was removed. Frequently arising artifacts following the same basic pattern, such as eye blinks or cardiac artifacts, were removed via SSP using projectors that were calculated separately for the cardiac and blink artifacts. All data were high-pass filtered with 1 Hz to remove movement-related low-frequency artifacts. Finally, the data were downsampled to 1000 Hz. All trials with artifacts were excluded from further analysis. For each trial, we extracted pre-stimulus activity that ranged from -8.0 seconds to -0.1 seconds, with 0 indicating the onset of the electric stimuli. This time window was chosen as it effectively excludes any post-response or pre-stimulus activity. Each channel's data were mean corrected by the average across the trial (-8.0s to -0.1s).

Cortically constrained source estimation was performed on these data at the participant level using each participant's anatomy. Using FreeSurfer (https://surfer.nmr.mgh.harvard.edu/, v.5.3.0), the participants' cortical surfaces were extracted from the individually acquired T1-weighted MRI scans (3T scanner and 1 $mm^3$ voxel size). The MRI was acquired after both MEG sessions had been recorded. We used the overlapping spheres method[2] with 306 spheres for the forward model and a linearly constrained minimum variance (LCMV) beamformer as the inverse model. The data covariance matrix for the LCMV beamformer was computed per measurement block of each participant, and the matrix was regularized by setting the values below the median to the median eigenvalue of the data covariance matrix. The noise covariance was obtained from a five-minute empty room recording on the same day as the measurement. A single shared source kernel was obtained for each measurement block of every participant.

For every trial and each participant, the source-reconstructed MEG data were projected to the default cortical anatomy (MNI 152 with 15,002 vertices) using registered spheres extracted using FreeSurfer and MRIs of individual participants[3] and then down-sampled to 250 Hz. We used the Mindboggle atlas[4] to spatially reduce the data dimensions. For each of the 42 cortical regions in the atlas, for every trial, we obtained the first principal component from the vertices' time series within that region. To correct for volume conduction in the signal, symmetric orthogonalization[5] was applied to these PCA (principal component analysis) time series. This resulted in 42 multivariate time series for each trial and each participant. The row vectors of this orthogonalized matrix, obtained for every trial, were z-scored. The entire pre-processing pipeline described in this paragraph was applied to all trials of every participant, irrespective of the response type on a trial.

Finally, to resolve sign ambiguity inherent in source-reconstructed MEG data across participants, a sign-flip correction procedure[6] was applied to the entire dataset, i.e., across all trials and participants. Subsequently, this dataset was then fed to the TDE-HMM pipeline described in the next section.

### HMM model fitting

Since we were interested in recovering phase-related networks, the TDE-HMM was fit directly on the time series obtained after the pre-processing steps described in the previous section, as opposed to its power envelope. This preserved the cross-covariance within and across the underlying time series of the cortical regions.

The HMM-MAR toolbox[6] was used for fitting the TDE-HMM. We opted for six different networks as a reasonable trade-off between the spectral quality of the results and their redundancy. The embedding took place separately in a 60 ms window (i.e., a 15-time point window for a sampling frequency of 250 Hz) for each trial. Since time embedding increases the number of rows of the data from 42 to 42 times the number of window samples, an additional PCA step was performed along the spatial dimension. The number of components retained was 84 (42 times 2) following our previous pipeline[7] and as recommended in reference 11. A full covariance matrix with an inverse Wishart prior was used to characterize each network. The network transition matrix of an HMM is described by a two-dimensional multinomial distribution. The conjugate prior for a two-dimensional multinomial distribution is a two-dimensional Dirichlet distribution. A two-dimensional Dirichlet distribution can be parameterized by concentration parameters alpha. The diagonal alphas were set to 10, and the off-diagonal were set to 1. The 'zero mean' option in the HMM toolbox was chosen to ensure that the mean of the time series did not take part in driving the networks. We used the stochastic version of variational inference for the HMM to speed up the fitting process. The 'HMM-MAR'-type initialization was used to start the optimization process (for details, see ref.[6]). A single HMM was fit across all participants and trials where the SOA was in the range critical SOA +- 10ms, irrespective of correct or incorrect stimuli detection. We hypothesized that correct and incorrect detections involve the same spontaneous networks, but their temporal interactions differ in the pre-stimulus epochs, leading to different detections.

### Data-driven frequency bands

We investigated the spectral connectivity patterns across the different networks. The objective was to uncover significant coherence within frequency bands in the respective networks. The HMM output includes a multidimensional "gamma" time series containing the probability for each network at a given point in time. A network was considered to be active at a given time point if the gamma value for that specific network was greater than or equal to 0.85. A contiguous block of time for which the gamma value for a specific network remains higher than 0.85 is referred to as a 'network visit'. These network time courses allowed the extraction of network-specific data for each trial for further analysis. For each HMM network, we filtered the network-specific data for all trials between 1 and 45 Hz. Then, we calculated the Fourier transform of the data using a multi-taper approach to extract the frequency components from the short segments of each network visit[8]. Seven Slepian tapers with a time-bandwidth product of 4 were used, resulting in a frequency resolution of 0.5 Hz. Subsequently, we calculated the coherence and power spectral density of these binned frequency domain data separately for every network within each trial. For each trial, the coherence and the power spectral density obtained were three-dimensional matrices of size f (number of frequency bins) by N (42 cortical locations) by N. We call these trial-level coherence matrices.

Based on the trial-level coherence matrices, we performed a frequency band-specific analysis. The lower triangular portion of the trial-level coherence matrix obtained above was vectorized across columns for each subject and all their trials. This resulted in 903 (lower triangular entries of a 42 by 42 matrix including the diagonal) by f (number of frequency bins) matrices for each trial. Subsequently, we averaged all matrices across trials and participants along the spectral dimension (number of frequency bins), resulting in a single 903 by f matrix per network. This procedure was repeated for all six networks. Finally, we concatenated matrices across all networks, resulting in a group-level coherence matrix (f by 903 by 6). We factorized the group-level coherence matrix into four frequency modes using a non-negative matrix factorization (NNMF)[9]. In contrast to the traditional definition of frequency bands in electrophysiological data, i.e., delta, theta, alpha, beta, and gamma, these frequency modes are data-driven descriptions of the spectral space and were obtained from non-negative matrix factorization of the Fourier-transformed data[8]. Canonical definitions of frequency bands assign equal weight to each frequency bin within a band for every subject, which might not be suitable for analyses across a large dataset.

The frequency mode weights are the NNMF weights obtained from the NNMF estimation (which, just like a regression coefficient, are unitless because coherence is unitless) (see supplementary figure 1). Three modes mostly correspond to the canonical alpha, beta, and high beta/gamma band, whereas the fourth represented the 1/f noise in neural signals. Since NNMF does not guarantee a unique solution, we performed multiple instances of the factorization. In practice, we could obtain frequency modes corresponding to the classical frequency bands within four algorithm iterations. At each instance, we visualized the output to ensure the frequency specificity of the frequency modes. The output stability was ensured using 'robust NNMF', a variant of the NNMF algorithm[8]. We then computed the inner product between the trial-and group level coherence matrix and the frequency modes obtained above. We called these the trial-level and group-level coherence projections, respectively. While these frequency modes were derived from coherence, they can be applied to power measures or any other frequency-specific measure. We used these factors to calculate the power spectra for each of the 42 regions and to obtain trial-level and group-level power projections.

### Diffusion map analysis

We used diffusion maps to recover a low-dimensional network embedding from high-dimensional cortical data[10,11]. The objective of the diffusion maps algorithm is to uncover the underlying structure of high-dimensional data, in this case, multivariate brain data. The following steps were performed to calculate network-specific embeddings: We masked the original dataset, i.e., the one without any delay embedding, using the network probability time course (HMM gamma output). The probability threshold for gamma was the same 0.85 as used to perform the multi-taper spectral analysis. The network time courses allowed the extraction of network-specific data for each trial for further analysis. Using the network-specific data, we computed a network-specific covariance matrix. The covariance matrix was then used to perform diffusion map analysis.

The diffusion map algorithm can be broken down into four main steps:

1. Define a similarity measure between pairs of data points
2. Construct a graph using the similarity measure
3. Compute the transition matrix and perform a diffusion process
4. Calculate the diffusion coordinates for a low-dimensional embedding

Step 1: Similarity measure

The similarity measure used in diffusion maps is typically a function that measures the distance between two data points. Let **C** be the covariance matrix for all brain regions, where **C (i, j)** represents the covariance between regions **i** and **j**.

To calculate the similarity between regions **i** and **j** based on the covariance matrix, we can use the following steps:

**C (i, :)** and **C (j, :)** are covariance vectors, i.e., the rows of the covariance matrix corresponding to regions **i** and **j**, respectively.

The similarity measure between **i** and **j** is defined as the Spearman rank correlation $\rho\big(C(i,:), C(j,:)\big)$

Then we can define a similarity matrix:

$$W(i,j) = \rho\big(C(i,:), C(j,:)\big)$$

Step 2: Graph construction

Using the similarity measure, we can construct a weighted graph G = (V, E) where V is the set of brain regions and E is the set of edges between brain regions. The weight of an edge between two brain regions i and j is given by the similarity measure calculated previously.

$$w_{ij} = W(i,j)$$

This similarity measure can be said to capture higher-order relationships between regions i and j that play an important role in complex systems such as the brain[12–14].

Step 3: Diffusion process

To perform the diffusion process, we compute the diffusion operator $P_\alpha$, which is defined as:

$P_\alpha = {D_\alpha}^{-1} W_\alpha$. Here, $\alpha \in [0,1]$ is the diffusion parameter used by the diffusion operator. The parameter α was set to 0.5 as per previous recommendation for large-scale brain data[15–17]. $W_\alpha$ and $D_\alpha$ are defined as follows:

$$W_\alpha = D^{-1/\alpha} W D^{-1/\alpha}$$

$D(i,i) = \sum_j W(i,j)$, with the off-diagonal elements set to 0

$$D_\alpha = \sum_j W_\alpha(i,j).$$

D is the degree matrix of the graph derived from W. Once $P_\alpha$ has been calculated, we can run the diffusion process multiple times by taking larger powers of $P_\alpha$, i.e., $P_\alpha^t$ , where t is called the time or the scale parameter[11]. We ran a single-step diffusion process with t=1. This was in line with the previous usage of diffusion maps for large-scale brain datasets[15–17].

Step 4: Diffusion coordinates and low-dimensional embedding

Once $P_{0.5}^1$ has been calculated, we can perform a generalized eigenvalue decomposition of the $P_{0.5}^1$ matrix.

$$P_{0.5(i,j)}^1 = \sum_l \lambda_l^1 \psi_l(x_i) \phi_l(x_j)$$

where $\{\lambda_l^1\}$ is the sequence of eigenvalues of $P_{0.5}^1$. $\{\psi_l\}$ and $\{\phi_l\}$ are the right and left eigenvectors, respectively.

Finally, the diffusion map can be defined as the new set of coordinates for the data that we can find using the right eigenvectors.

$$\psi_1(x) = \left(\lambda_1^1 \psi_1(x), \lambda_2^1 \psi_2(x), \ldots \lambda_k^1 \psi_k(x)\right)$$

Using $\psi_1(x)$ we can embed the data into an Euclidean space, which we term as the connectivity manifold in our results. For a complete theoretical description of diffusion maps, please refer to reference[9].

The dimensionality was reduced from 42 to 5. Only the top three dimensions were used for subsequent analysis by looking at the elbow of the scree plot. The top three dimensions captured 75% of the variance in the data. The above procedure was implemented using custom-written scripts and Brainspace toolbox[18].

### Figures

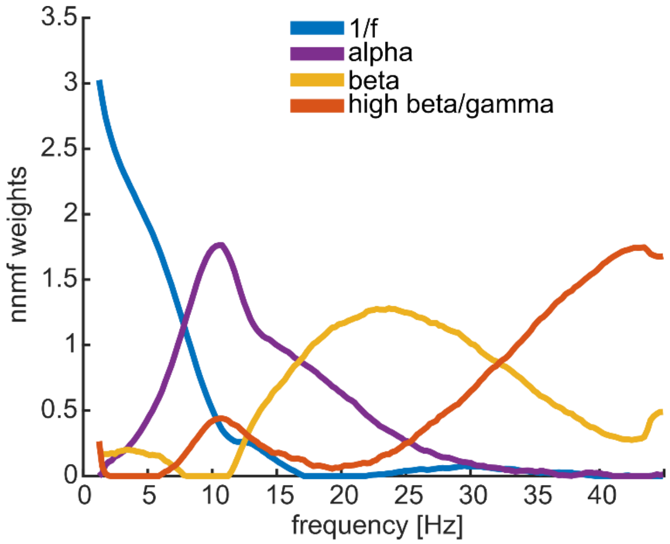


**Supplementary figure 1: Data-driven frequency bands, referred to as frequency modes in the text.** The x-axis represents frequency in Hz and the y-axis represents the weights obtained from the non-negative matrix factorisation (NNMF) in arbitrary units. The frequency resolution of the modes is 0.5 Hz.

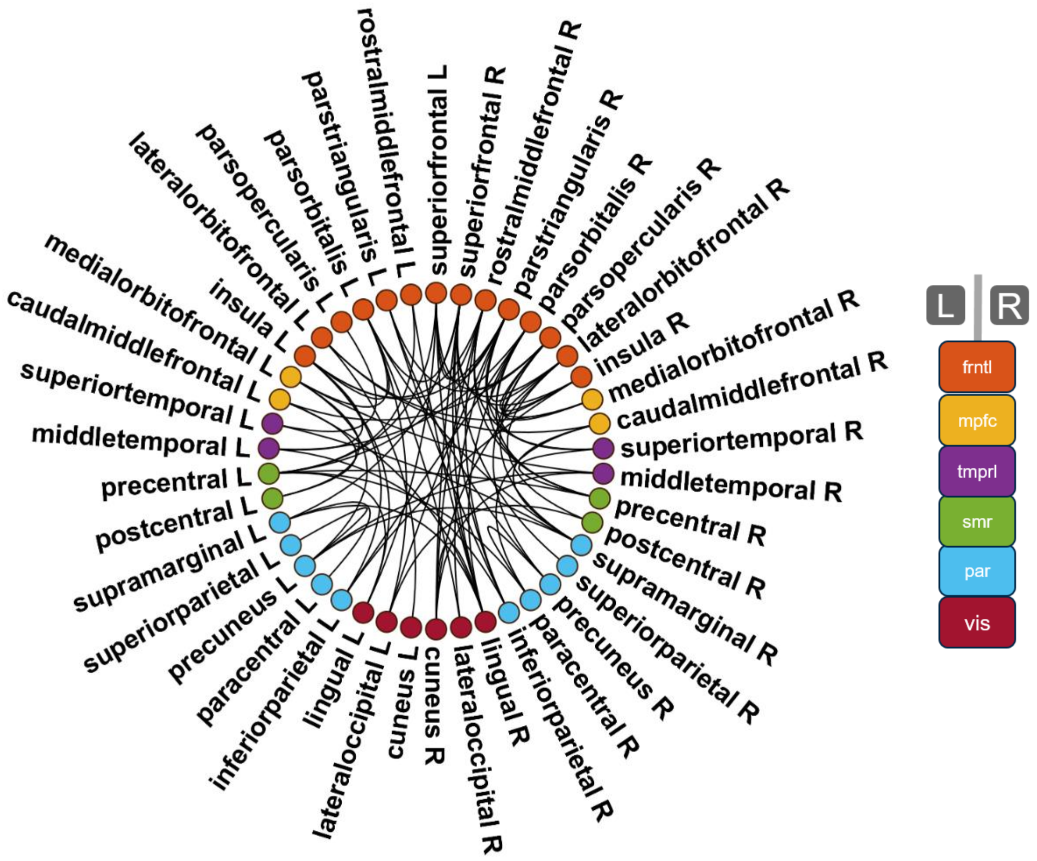


**Supplementary figure 2: Brain region labels for the ring figures.** The brain regions are based on the Mindboggle atlas.

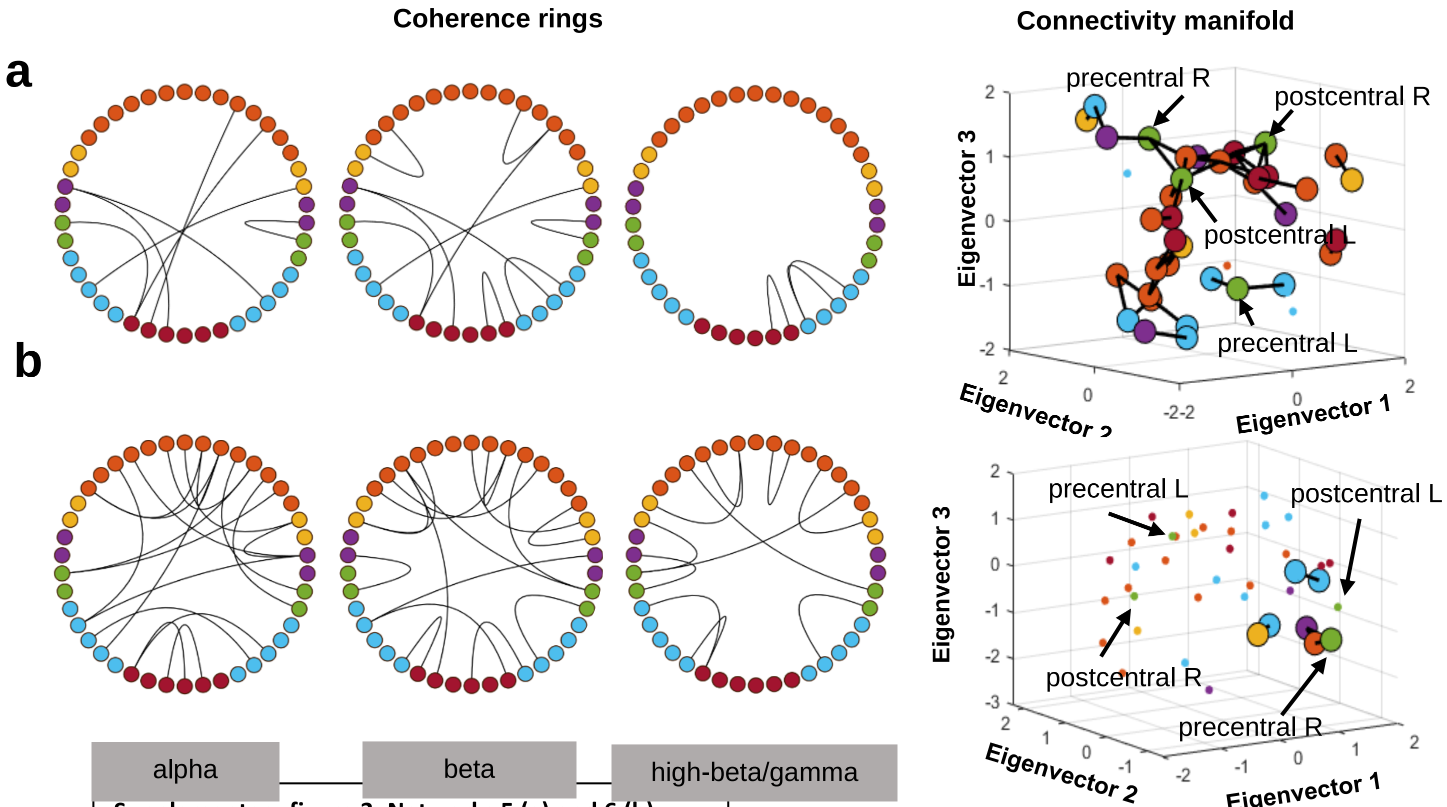


**Supplementary figure 3: Networks 5 (a) and 6 (b).**
For details on the legend see main figure 2.

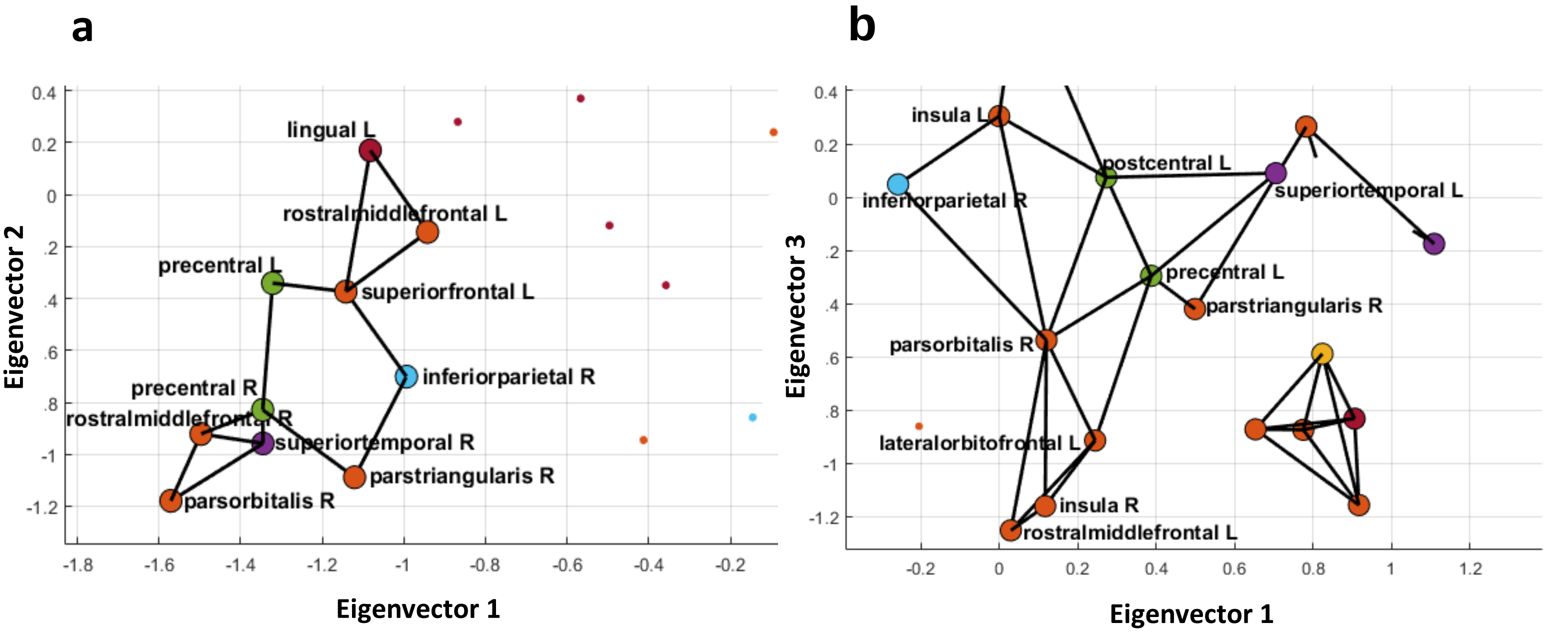


**Supplementary figure 4: Zoomed version of the connectivity manifold**. **a)** Cross-brain network **b)** High-beta/gamma network. In both the figures the inferior parietal R is connected to sensory motor regions via two frontal areas.

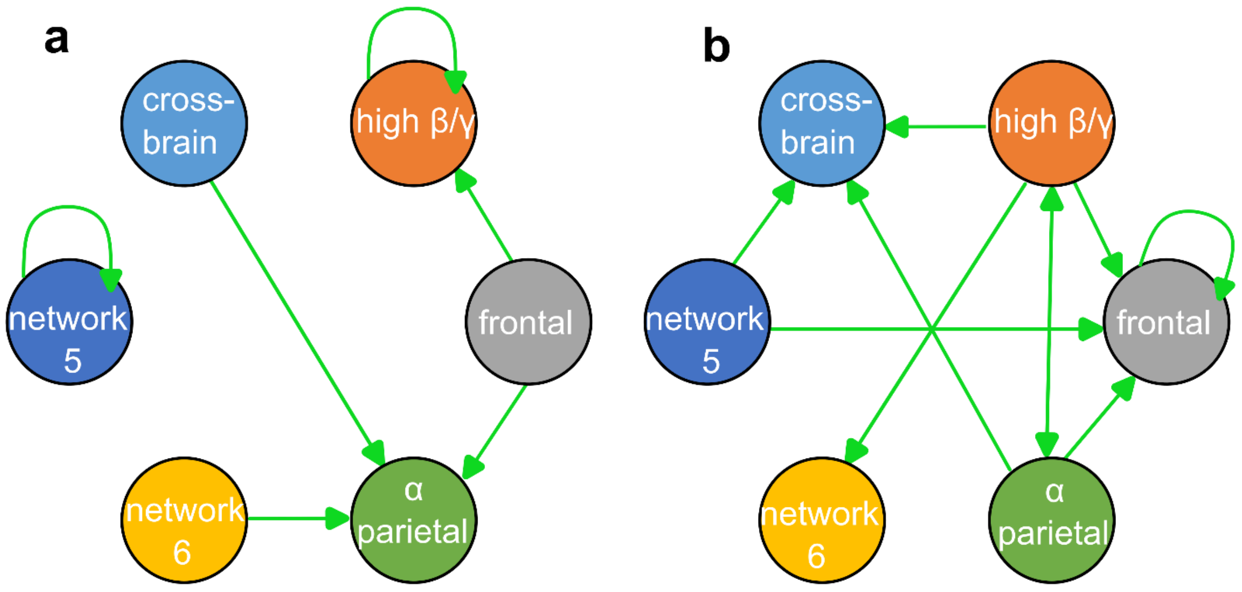


**Supplementary figure 5: Complete set of network transitions**. **a)** Significantly increased network transition probabilities when participants incorrectly detected the stimuli. **b)** Significantly increased network transition probabilities when participants correctly detected the stimuli, $p<0.05$. Not depicted transitions were statistically equally likely across perceptions.

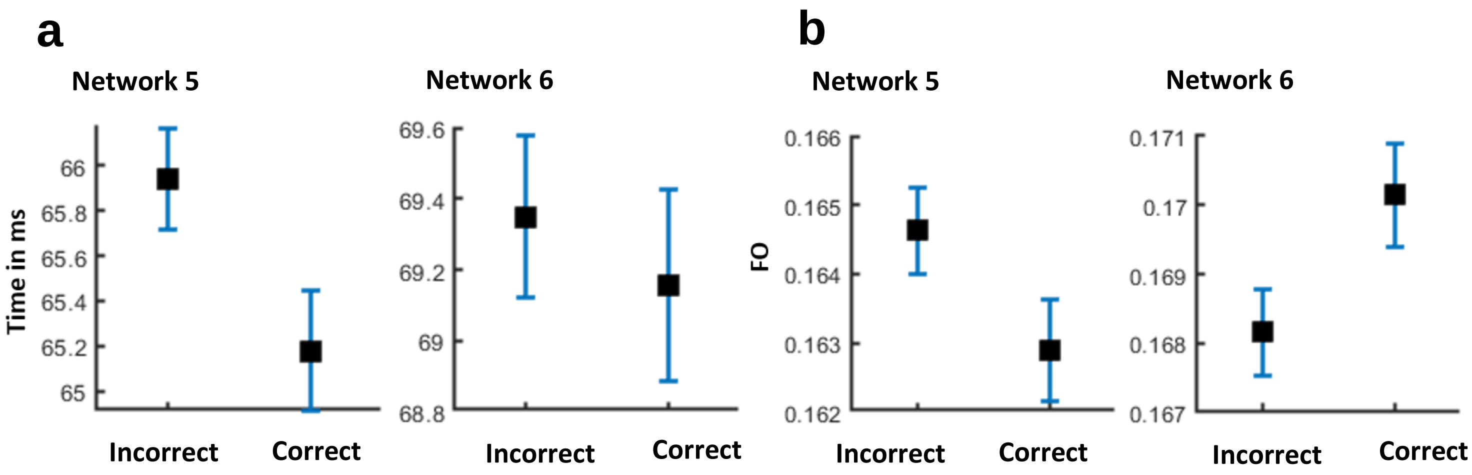


**Supplementary figure 6: Temporal properties of networks 5 and 6.**
**a)** Lifetime **b)** Fractional occupancy (FO)

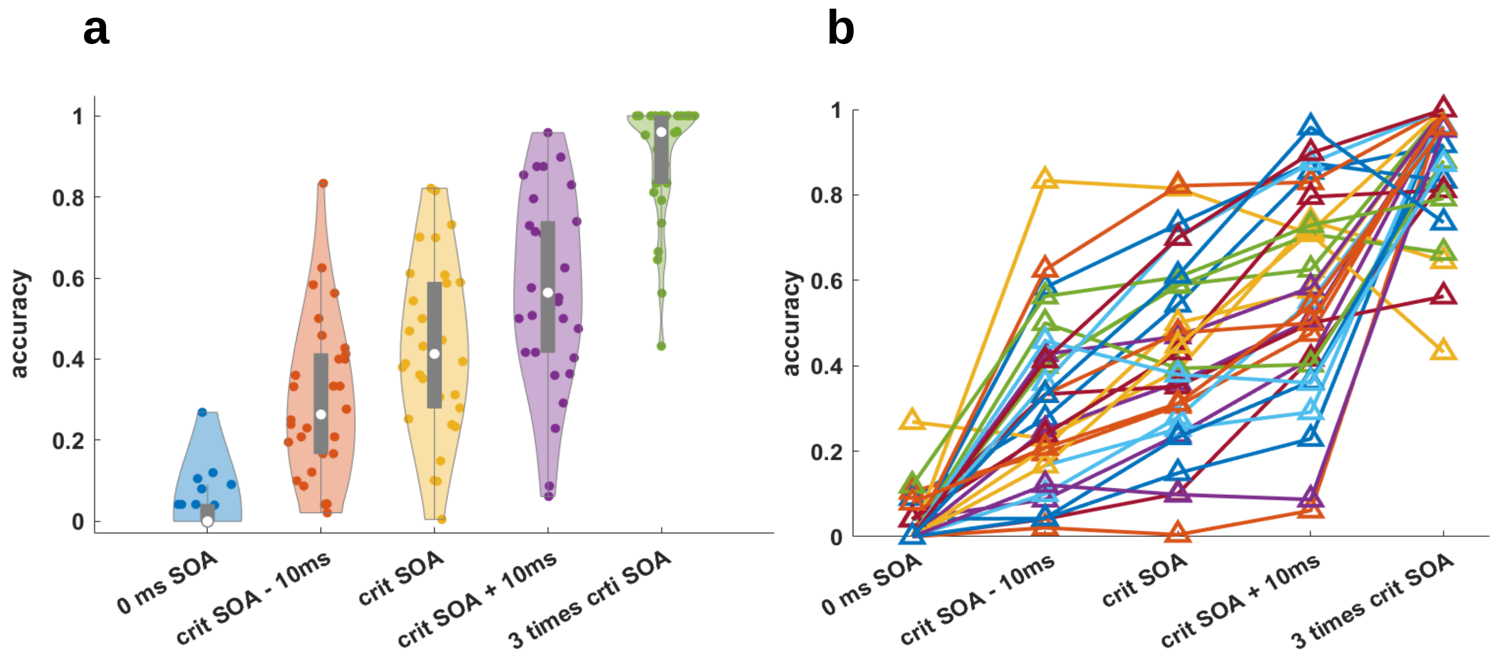


**Supplementary figure 7: Accuracy plots.**
Accuracy plot under different SOA (stimulus onset asynchrony) conditions. crit SOA: critical SOA. Accuracy was calculated per subject and per SOA condition. Accuracy was calculated by dividing the number of trials for which participants detected two electric pulses by the total number of trials.
**a)** Each violin plot includes a boxplot (grey). Box edges are formed by the first 25th percentile quartile to third 75th percentile quartile. White dots are the medians. Data minima/maxima control the whisker length. Individual scatter points within a violin plot are individual participant accuracies in a specific SOA condition. The width of each violin plot provides a probability density estimate for the accuracy given the participant data.
**b)** Individual participant accuracy plots under different SOA (stimulus onset asynchrony) conditions. Each triangle represents an individual participant's accuracy under an SOA condition. For a given participant, triangles under different SOA conditions are connected using a line plot.

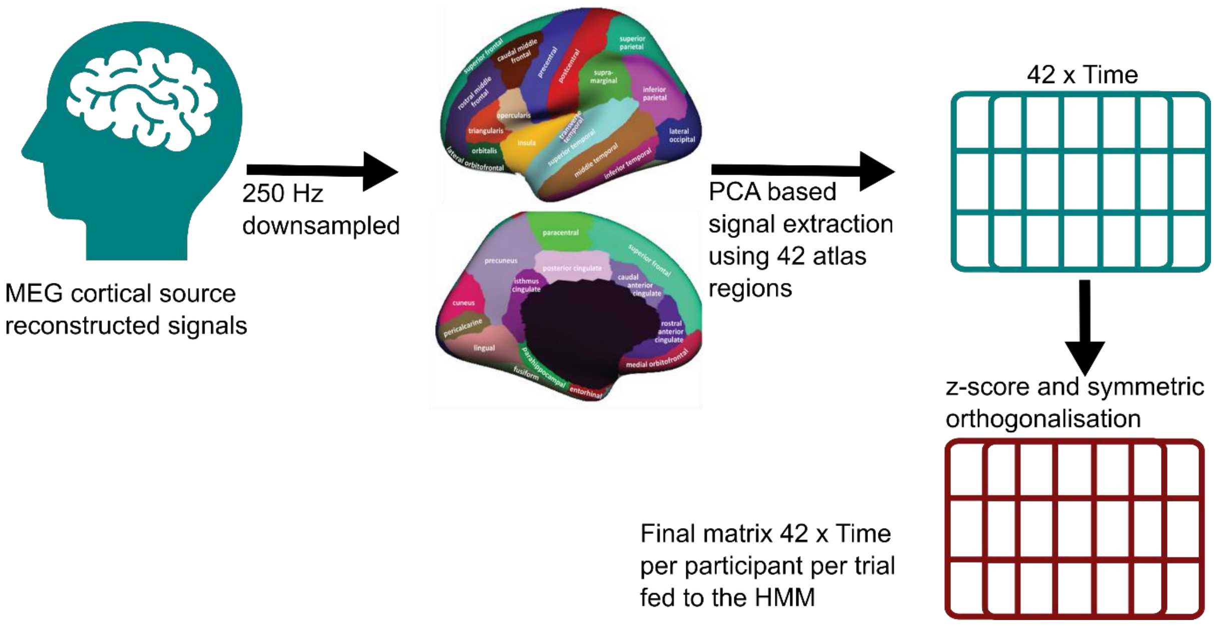


**Supplementary figure 8: A summary of the data processing pipeline for individual trials**.
